# Whole-Genome Sequencing on 220 Alfalfa (Medicago sativa L.) Accessions Identified Association of DREB1C Gene with Fall Dormancy Height

**DOI:** 10.1101/2021.03.29.437533

**Authors:** Fan Zhang, Junmei Kang, Ruicai Long, Mingna Li, Yan Sun, Zhen Wang, Zhiwu Zhang, Qingchuan Yang

**Affiliations:** Institute of Animal Science, Chinese Academy of Agricultural Sciences, Beijing, China; Department of Crop and Soil Sciences, Washington State University, Pullman, WA, USA; Department of Turf Science and Engineering, College of Grassland Science and Technology, China Agricultural University, Beijing, China

**Keywords:** GWAS, fall dormancy, Medicago sativa, DREB1C, freezing tolerance

## Abstract

Fall dormancy (FD) is one of the most important traits of alfalfa (Medicago sativa) for cultivar selection to overcome winter stress. Although transcriptomics, proteomics analysis, and QTL mapping have revealed some important genes correlated with FD, the genetic architecture of this trait is still unclear. There are no applicable genes or markers for selection, which hinders progress in the genetic research and molecular breeding for the trait. We conducted whole-genome sequencing (WGS) on 220 alfalfa accessions at 10x depth. Among the 875,023 SNPs, four of them were associated with FD height using GWAS. One SNP located on chromosome 6 is in linkage disequilibrium with dehydration-responsive element-binding protein 1C (DREB1C). Furthermore, seven DREB genes are clustered in this region, one of which has previously been shown to enhance freezing tolerance in the model plant Medicago truncatula. The candidate genes uncovered by our research will benefit the transgenic and CRISPR-Cas9 research of FD in alfalfa. This gene will also be useful for molecular marker development and marker-associated breeding of FD for alfalfa.

## Introduction

Fall dormancy (FD) is the adaptive growth characteristic of alfalfa in autumn and can be evaluated by the regrowth shoot height at 25-30 days after final cutting (Teuber et al. 1998). The taller alfalfa grows in the fall, the less dormant it is. The FD level of alfalfa varieties can be classified from 1 to 11, where 1 represents most dormant and 11 represents least dormant (Teuber et al. 1998). Dormancy is induced by falling temperature and shortening photoperiod. Dormancy can improve plant survival in winter and increase freezing tolerance. FD alfalfa cultivars have better winter survival and hardiness than non-dormant alfalfa cultivars (Barnes et al. 1979; Smith 1961; STOUT 1985; STOUT and HALL 1989). FD level can be considered one of the indices for alfalfa cultivar selection in specific regions. There is a trade-off between FD and yield. High FD could increase overwintering ability, but decrease the potential yield (Avci et al. 2018; STOUT and HALL 1989). Breaking down the linkage between FD and yield from genetic level could benefit for both yield and winter hardiness. Studying the genetic architecture of FD could be used to improve those high yield alfalfa cultivars with bad cold tolerance (Li et al. 2015; Munjal et al. 2018).

The genetic mechanism of FD in alfalfa has been studied using genomics, transcriptomics, and proteomics. QTL mapping between different FD level alfalfa germplasms has been conducted in several studies. Seventy-one QTL related to FD traits have been identified, and 15 of them can be detected in more than one environment (Li et al. 2015). Another study identified 45 significantly associated QTL for FD (Adhikari et al. 2018). Both of these groups found that FD and winter hardiness have independent inheritance and could be improved independently. A gene expression related experiment showed that a cold acclimation responsive gene, RootCAR1, was positively associated with alfalfa winter survival (Cunningham et al. 2001). More FD-related genes have been found using transcriptomics technology. Forty-four genes were identified by comparison of the transcriptomes from non-FD versus FD cultivars’ leaves. The transcription of the genes IAA-amino acid hydrolase ILR1-like 1, abscisic acid receptor PYL8, and monogalactosyldiacylglycerol synthase-3 were suggested to be involved in regulating FD in alfalfa (Du et al. 2017). Eight significantly differentially expressed transcription factors related to CBF and ABRE-BFs were identified (Liu et al. 2019). Furthermore, FD-related proteins were analyzed using proteomics and metabolomics. A total of 90 proteins were different between FD and non-FD alfalfa. One of these was thiazole biosynthetic enzyme (MsThi), which is important for alfalfa growth (Du et al. 2018). Raffinose family oligosaccharide (RFO) metabolism was shown to be involved in short photoperiod-induced freezing tolerance in dormant alfalfa cultivars (Bertrand et al. 2017).

Genome-wide association studies (GWAS) are widely used to locate precise SNPs associated with phenotypes using historical recombination information. With the development of the sequencing method and genetic analysis method of GWAS, the candidate SNP markers for important quantitative traits have been identified across different populations (Biazzi et al. 2017; Wang et al. 2016; Yu et al. 2016). Based on 198 accessions, Longxi-Yu reported a set of GWAS results, including 19 SNPs associated with drought resistance (Zhang et al. 2015). 36 SNPs associated with salt tolerance (Yu et al. 2016), 42 associated with plant growth and forage production (Liu and Yu 2017), and 131 markers associated with 26 forage quality traits (Lin et al. 2020). Using the 336 genotypes, Wangzan conducted GWAS for several key alfalfa traits, including fiber-related traits and digestibility (Wang et al. 2016), crude protein and mineral concentrations (Jia et al. 2017), and nine biomass-related traits (Wang et al. 2020). Another study reported several SNPs associated with forage quality using half-sib progeny developed from three cultivars (Biazzi et al. 2017). Furthermore, several genes were revealed to be related to some important traits of alfalfa. For example, several stress-responsive genes associated with yield under water deficit were identified, including leucine-rich repeat receptor-like kinase, B3 DNA-binding domain protein, translation initiation factor IF2, and phospholipase-like protein (Liu and Yu 2017). Two markers linked to NIR-NBS-LRR genes were significantly associated with Verticillium wilt resistance (Yu et al. 2017). A cell wall biosynthesis gene associated with several cell wall–related traits that affect alfalfa nutritive value has been identified (Sakiroglu and Brummer 2017). MsACR11 has an effect on plant height, which could significantly increase the height of transgenic arabidopsis (Wang et al. 2020).

Although many GWAS have been performed in alfalfa, few of these have focused on FD. Furthermore, with the lack of reference genome in alfalfa, most previous studies have used the Medicago truncatula genome as a reference (Sakiroglu and Brummer 2017; Wang et al. 2020; Yu et al. 2017). With the technological development of the genome assembly, the genome of the cultivar “Zhongmu No.1’’ has been released. A flowering locus T homolog gene, MsFTa2, may be associated with fall dormancy and salt resistance and was identified using the GWAS method and the new reference genome for alfalfa (Shen et al. 2020).

In this study, we evaluated FD height traits in 220 accessions and conducted WGS with the mean depth of 10x sequencing for every accession. A total of 875,023 SNPs were detected in the alfalfa reference genome and used for conducting GWAS. The objectives of this experiment were (1) evaluate the linkage disequilibrium (LD) and population structure of alfalfa, (2) identify SNPs associated with FD, and (3) infer the candidate genes controlling FD.

## Results

### Phenotypic variance

To identify the genetic difference in FD height for different seeds, the variance of the phenotype was analyzed. The effect of accession (genotype), year, individual, and accessions– year interaction were estimated (Table1). FD height was recorded for two years: 2018 and 2019. The variance of FD height in 220 accessions is given in Table 1. Significant differences in accession, year, and accession-year interaction were observed with p-values smaller than 0.0001. Slightly significant differences among replications were also observed (p=0.02), while the variation of individuals was not significant (p=0.12), indicating that the difference in FD between individuals is a random effect (Table 1). The broad-sense heritability was 69.2% for FD height.

**Table 1.** Analysis of variance for FD height in GWAS population over two years.

| Source | df | Type III SS | F-value | P-value | Variance |
| --- | --- | --- | --- | --- | --- |
| accession | 219 | 111676.00 | 10.73 | <.0001 | 15.41 |
| year | 1 | 116825.50 | 2458.49 | <.0001 | 35.39 |
| ind(accession) | 866 | 43915.99 | 1.07 | 0.12 | 0.53 |
| repeat | 2 | 382.01 | 4.02 | 0.02 | 0.07 |
| accession*year | 204 | 22669.18 | 2.34 | <.0001 | 4.24 |

The effect of FD height on yield-related traits was estimated. Different correlation coefficients were observed between FD height and other yield-related traits. The highest correlation coefficient (0.65, p<0.001) was found between FD height and regrowth. The plant height also showed a significant correlation (p<0.001) with FD height, and the correlation coefficient is 0.51 for all three-cut alfalfa. Leaf length and leaf width were significantly correlated with FD height at a relatively smaller correlation coefficient (p<0.001). Spring vigor and three-cut yields are correlated with FD height at the 0.01 significance level. The correlation between FD height and flowering time is not significant (Figure 1). Taken together, these results indicate that FD height is significantly associated with yield and yield-related traits (regrowth, plant height, leaf length, leaf width, and spring vigor).

**Figure 1.**
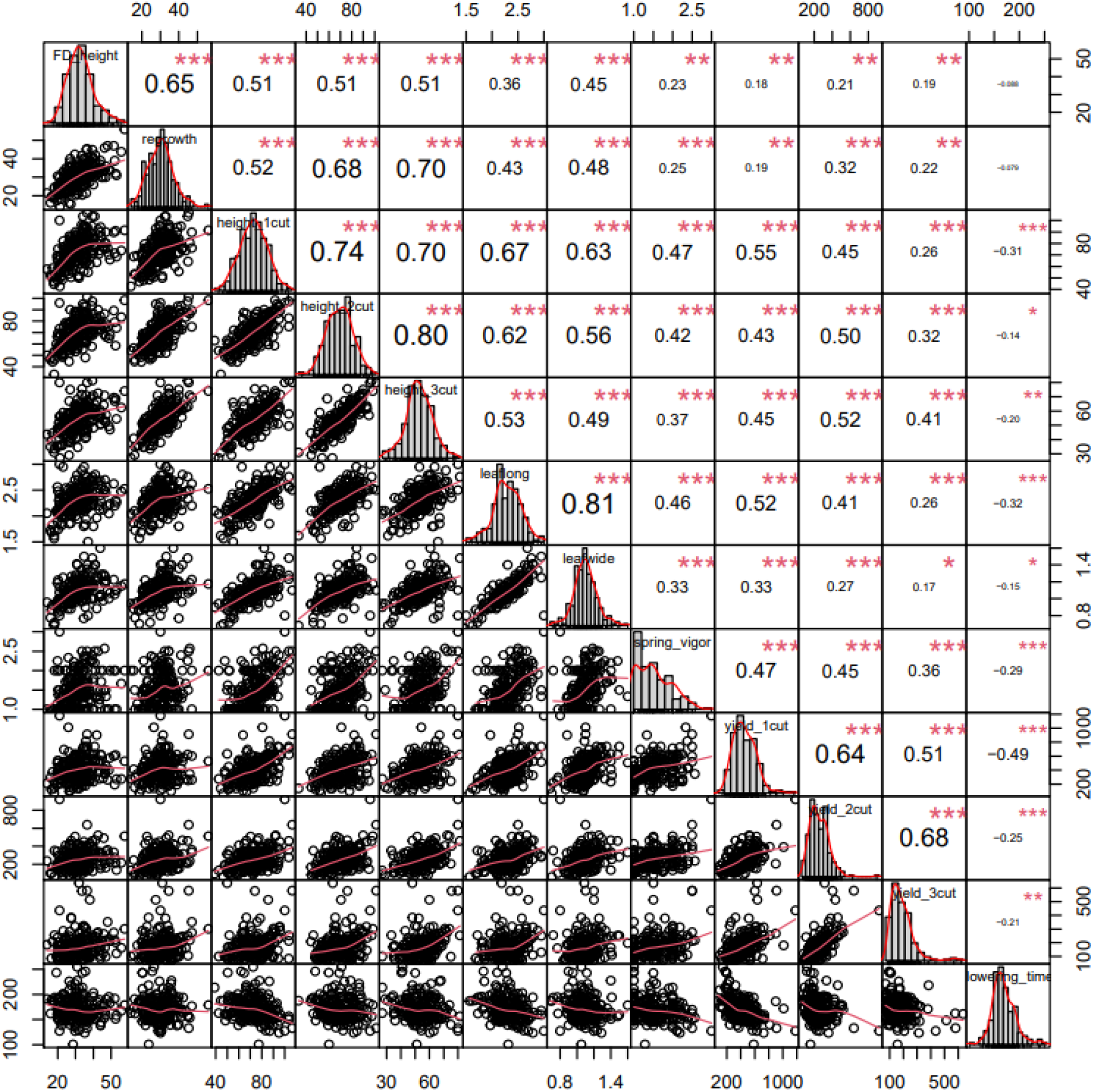
Correlations and distributions of 2019 FD height with 2020 yield-related traits. The histograms on the diagonal are the distributions for each trait. The upper triangle consists of the correlation coefficients among traits. The lower triangle consists of pairwise scatter plots. The signal *, **, and *** represent 0.05, 0.01 and 0.001 significant levels, respectively.

### Genotype calling and allele frequencies

A total of 32.2 million SNPs were detected using the BWA-SAMTools-VarScan pipeline. After filtering the SNPs with a missing value >10%, minimum mean read depth >20, and minor allele frequency (MAF) >0.05, there were 875,023 SNPs that passed the threshold and were confirmed for further analysis. Among these 875,023 SNPs, the average SNP density in the genome was one SNP per 26.95 bp (among the 814,517 SNP with SNP interval less than 300 bp) (Figure 2A, C). There were 60,506 SNP with an interval larger than 300bp, and the average SNP interval for these was 11128.66 bp (Figure 2C). The distribution of the MAF value showed that the value equal to 0.06 has the highest density (Figure 2B, detail data not shown). The SNP number range is similar for MAF values from 0.2 to 0.5 (Figure 2B). These results demonstrate that the markers can be assigned to unbiased chromosomes and distinct positions.

**Figure 2.**
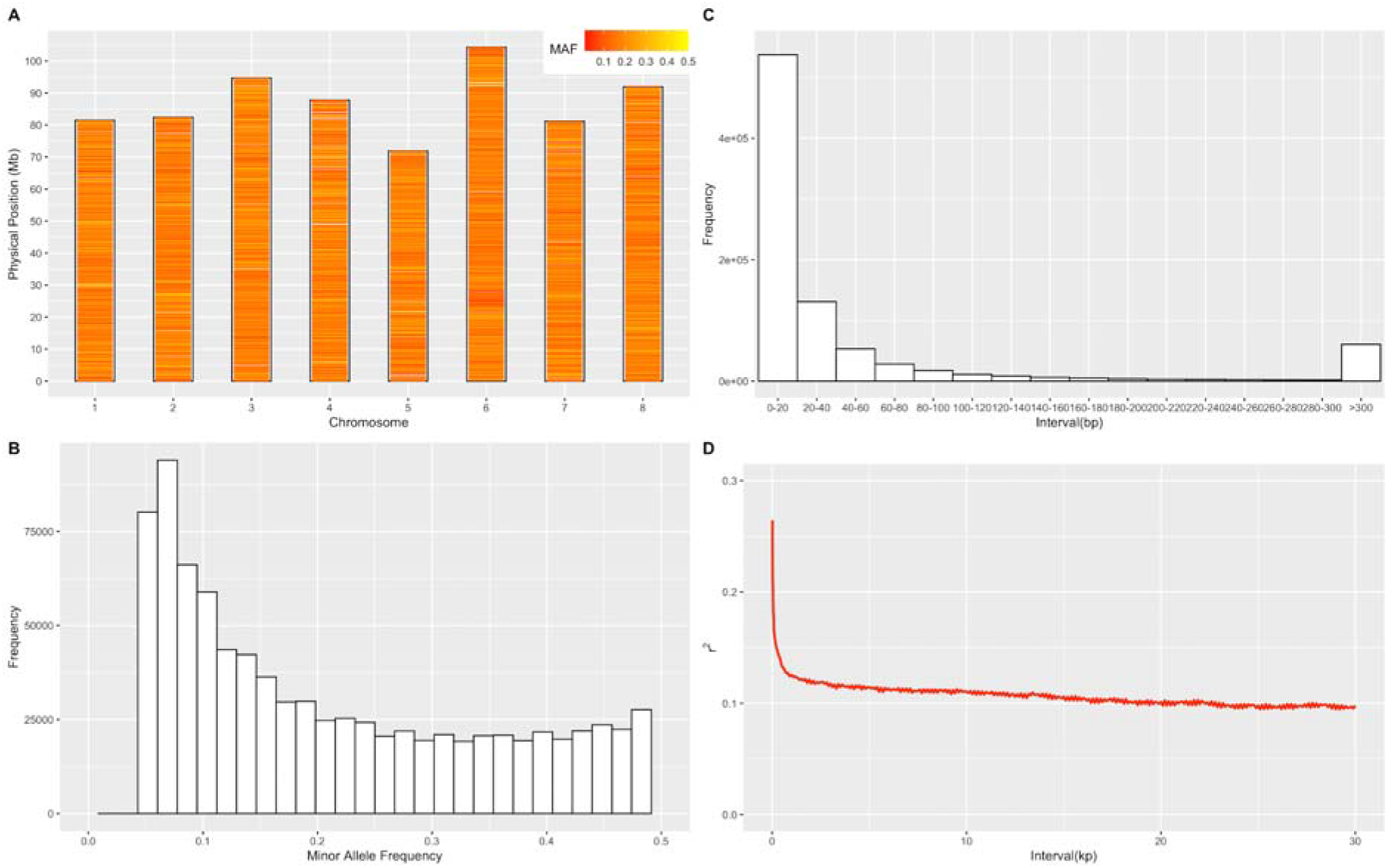
Properties of single nucleotide polymorphisms (SNPs). In total, 220 alfalfa accessions were genotyped by WGS; 875,023 SNPs passed filters and quality control. Marker distributions are displayed as a heatmap on 8 chromosomes by minor allele frequency (MAF) (A). MAF distribution is shown in a histogram (B). Marker density is displayed by histogram according to the interval of adjacent SNPs (C). LD decay is shown according to the red trend line (D).

### Population structure

The accessions used in our research originated from different countries, especially the USDA-GRIN collection, representing wide coverage of genetic diversity on alfalfa. To analyze population structure, the genome-wide SNP data generated by WGS was used for principal component analysis (PCA). The PCA scatter-plots showed a weak population structure for the 220 accessions consisting of different geographical origins with no clear subpopulation (Figure 3). In the scatterplot of PC1 and PC2 (Figure 3A), most accessions are mixed. But some accessions coming from China can be separated from those with other origins (red color). Similarly, some accessions from South America can be separated from the others (cyan color). The accessions from Europe, North America, and other Asian countries have wide and mixed distribution. The accessions from Africa also mixed with other accessions with a relatively low genetic diversity (green color) (Figure 3A-D). The first three PCs explained 2.6%, 1.8%, and 1.2% genetic variance separately; the total genetic variance explained by the first three PCs is 5.6%. These mixed clusters indicate that, although accessions come from different geographical origins, they still have a high degree of genetic relatedness.

**Figure 3.**
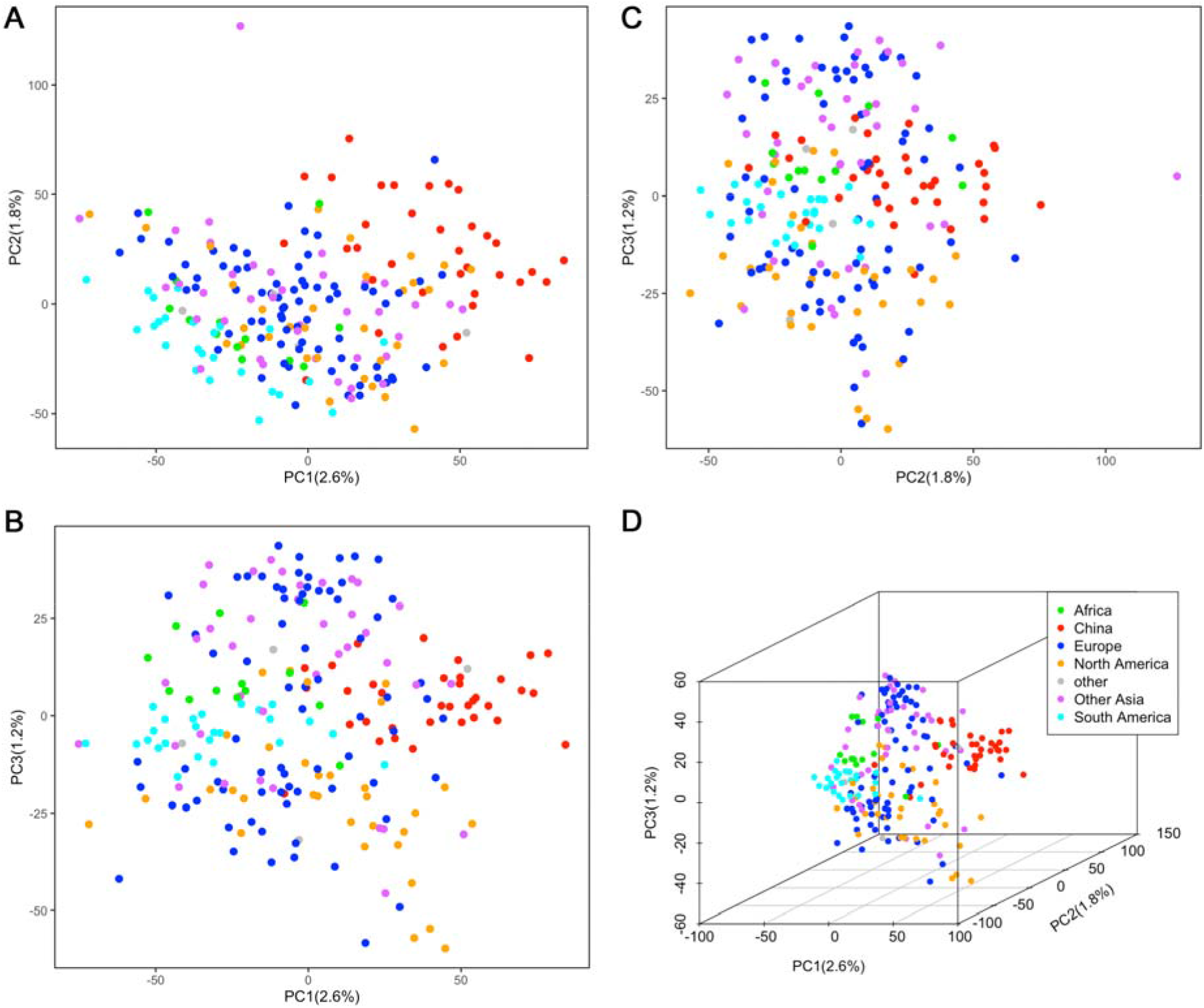
Population structure from the principal component analysis. A total of 875,023 SNPs and 220 alfalfa accessions were used to perform the principal component analysis. Population structure is shown as pairwise scatter plots (A, B, and C) and a 3D plot (D) of the first three principal components (PC) with colored points that define the seven groups. There are 13, 35, 73, 32, 4, 38, and 25 accessions in the groups corresponding to Africa, China, Europe, North America, other (not belonging to any other group), other Asia (except China), and South America, respectively.

### Linkage disequilibrium

To estimate the confidence interval of GWAS results, LD was analyzed using a total of 875,023 markers. The r2 of LD across all chromosomes was calculated and is presented in Figure 2D. A rapid drop in r2 was observed with the increase in physical distance. Over all the accessions, the value of r2 decreased quickly within 2 kb physical distance, then decreased slowly afterward (Figure 2D). The value of r2 decreased to 0.1 after 18 kb across all chromosomes. The results showed that extensive recombination took place during alfalfa evolution and distribution.

### GWAS and candidate genes for FD height

To identify SNP markers associated with FD height, genotypic and phenotypic data were analyzed using the Bayesian-information and Linkage-disequilibrium Iteratively Nested Keyway (BLINK) method in the BLINKC version software.

The quantile-quantile (Q-Q) plot results of marker-trait associations for FD height were illustrated using observed versus expected p-values (Figure 4). Any deviation from the expected red line implies SNP association with FD height. In the present study, a total of 4 SNPs passed the 1% threshold after a Bonferroni correction (p<1.14×10-8) and were associated with FD height (Figure 4, Table 2). Among them, two markers, chr2__23380918 and chr2__80663885, were located on chromosome 2. In addition, 2 markers (chr6__740339 and SNP chr6__69737395) were located on chromosome 6. The boxplot of four SNP genotypes associated with FD height is shown in Figure 5. The FD height difference between different genotypes can be observed in the four SNP boxplots. There is no homozygous genotype CC for SNP chr6_740339.

**Figure 4:**
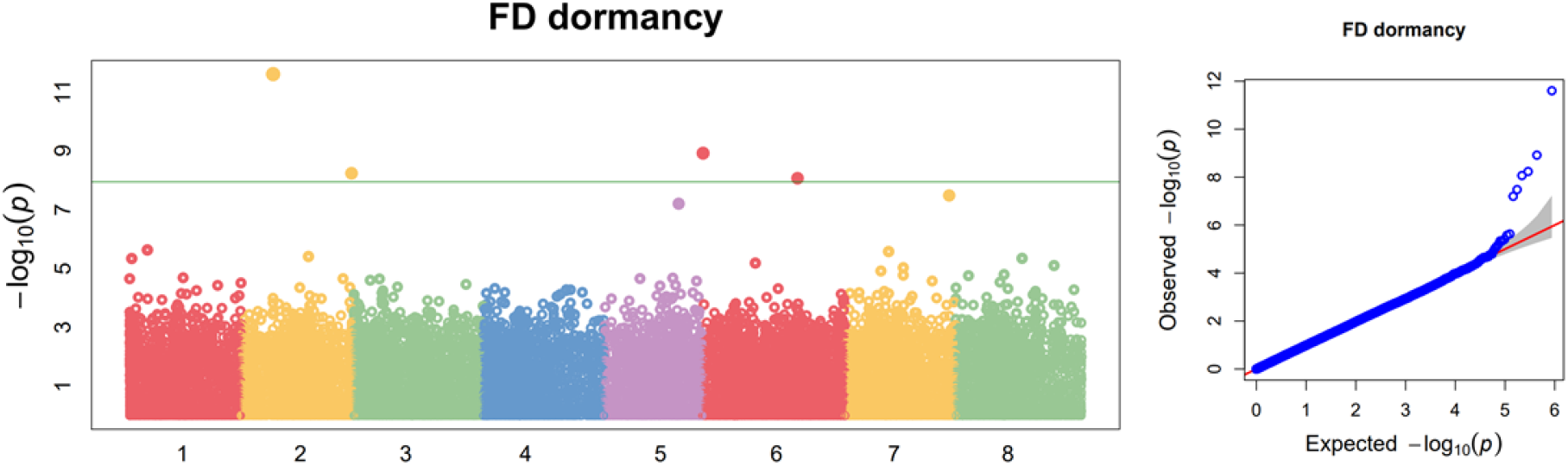
Manhattan and Quantile-Quantile (Q-Q) plot of fall dormancy height. The genome-wide association study was performed by BLINK C version software, with a significant p-value threshold set at P=0.01/875,023= 1.14×10-8(red line). We identified 4 significantly associated SNPs, which are shown in the Manhattan plot. Q-Q plots are displayed on the right panel. The Manhattan and Q-Q plots are generated using GAPIT3.

**Table 2.** Significant SNP markers associated with FD height.

| SNP | Allele | Chr | P-value | Candidate Gene | Gene Annotation |
| --- | --- | --- | --- | --- | --- |
| chr2__23380918 | C/T | 2 | 2.49E-12 | -- | -- |
| chr2__80663885 | C/A | 2 | 5.82E-09 | -- | -- |
| chr6__740339 | A/C | 6 | 1.21E-09 | Msa0867660 | signal anchor |
| chr6__69737395 | T/C | 6 | 8.59E-09 | Msa0892030 | DREB1C |

**Figure 5.**
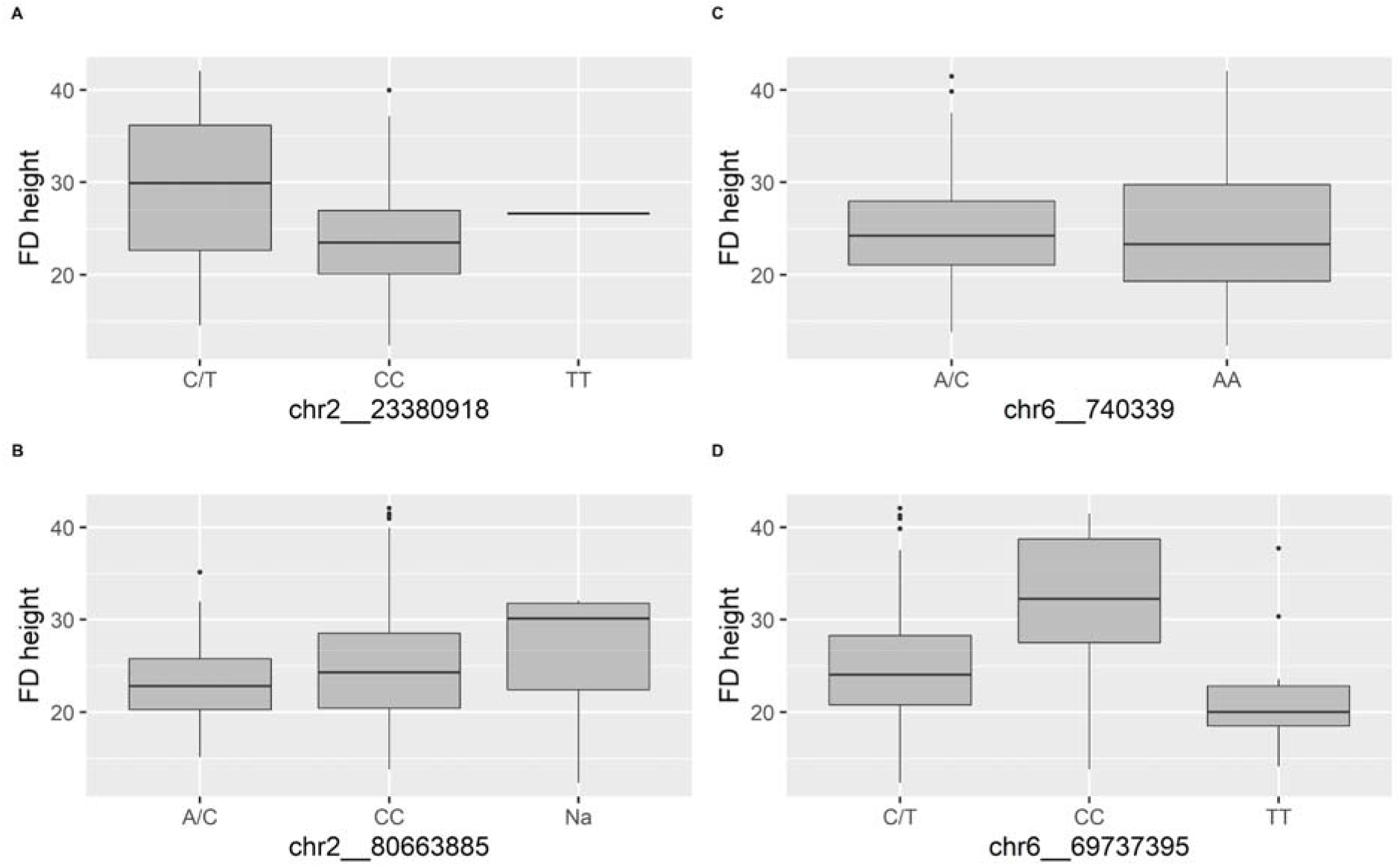
Distribution of FD height among the genotypes of the associated SNPs. There is only one individual with TT genotype for SNP chr2_23380918 and no individual with genotype of CC on SNP chr6_740339.

To identify potential candidate genes linked to SNP associated with FD height, candidate genes were screened based on the up- and downstream 18 kb LD interval information. Of the 4 significantly associated SNP, 2 of them are linked to known genes in the alfalfa genome (Table 2). There is one candidate gene (Msa0867660) with the annotated function of signal anchor protein connected to chr6__740339. Only one gene (Msa0892030), dehydration responsive element binding protein 1C (DREB1C), was connected with the SNP chr6__69737395 in the 36 kb genomic interval based on LD information (Supplemental Figure 1). We further checked the genes near SNP chr6__69737395 with a larger scope (500 kb interval up- and downstream).

Another six DREB genes were found upstream of this SNP (chr6:69234712-69716598). One of the six genes (Msa0891950) has been validated to be associated with freezing tolerance in *Medicago truncatula* (Supplemental Figure 2) (Chen et al. 2010). The transgenic Medicago truncatula has lower plant height than normal plants and has better tolerance for freezing. We further checked all DREB genes on chromosome 6, and found that there were nine in total (Supplemental Table 1). Another two DREB genes were Msa0893350 and Msa0893360 (chr6:76949157-76957350). There is only one cluster region with seven DREB genes on chromosome 6, and this region corresponds to our significant GWAS signal.

## Discussion

### The FD height variance of alfalfa

Because of the outcrossing and self-incompatibility feature of alfalfa, the genotype between different individuals is different. Most studies collect phenotypes using cloned plants (Li et al. 2015; Liu and Yu 2017; Sakiroglu and Brummer 2017; Wang et al. 2016). For one study, seed plants of the same accession were used for phenotype collection (Yu et al. 2016). Compared with cloned plants, the advantages of using seed plants are that they are easier to establish and have better accession representation. Furthermore, some experiments and assessment of phenotypes cause damage to the plants, such as winter survival, disease resistance, and root-related phenotypes. It is easier to rebuild the population using seed than by cloning plants. We checked the FD height variance of individuals based on our experiment. To check the influence of individuals’ genetic differences for FD height among the same accessions, we nested individuals in accession. Our results showed that the variance of individuals is not significant, while the variance of accessions is significant (Table 1). That means the influence of genetic differences between individuals among the same accession is not significant. The FD height variance can be considered as the genetic difference among accessions. Furthermore, our GWAS results showed that FD height has significantly associated SNPs and candidate genes. That means our FD height phenotype data collection strategy is feasible for GWAS. As for the GWAS power of other alfalfa agronomic traits, further experiments are needed, because the heritability and genetic differentiation level vary for different traits.

### The genetic diversity of alfalfa

The improvement of alfalfa is mainly based on outcrossing, and the genetic exchange of alfalfa always exists in alfalfa-growing regions (Li and Brummer 2012). To this end, nine distinct germplasms with global backgrounds were introduced in North America to cultivate new varieties (Barnes 1977). It is more difficult to distinguish alfalfa accessions based on geographic origin than for self-pollinating plants, such as soybean (Zhou et al. 2015) and rice (Huang et al. 2012). Also, diploid alfalfa (M. sativa subsp. caerulea and M. sativa subsp. falcata) can be clustered based on subspecies and geography (Sakiroglu and Brummer 2017; Şakiroğlu et al. 2010); the genetic diversity of tetraploid alfalfa (cultivated alfalfa) is more complicated. Based on our PCA results, the genetic differentiation among different origins is not very high, except the ones from China and South America.

The introduction of alfalfa to South America took place around the sixteenth century, while it was introduced to China 2000 years ago (Annicchiarico et al. 2015). Differentiation has occurred along with cultivation in the local region. This phenomenon has also been identified in previous studies (Qiang et al. 2015; Shen et al. 2020; Wang et al. 2020). The accessions coming from Europe and other Asian regions (regions except for China) are mixed and have a wide range. These accessions have the highest genetic diversity among all accessions with different geography. This result may be the reason that alfalfa originated from the Caucasus region: northeastern Turkey, Turkmenistan, and northwestern Iran (Michaud et al. 1988). This theory can also be strengthened by the knowledge that accession genetic diversity from North America and Africa is relatively low compared to European accessions. The cultivated alfalfa was exported to North America and Africa only around 200~300 years ago (Prosperi et al. 2014). This relatively short cultivation period also partially explains the low genetic diversity.

### The linkage disequilibrium of alfalfa

Our results also showed that the LD distance is very short (R^2 reduced to 0.1 after 18 kb). This is different from previous studies (Sakiroglu and Brummer 2017) and (Li et al. 2014). One major reason may be because we used a large number of SNPs (around 58 times more than the previous report). With increased marker density, it is easier to estimate historical recombination events. Another study showed that LD can be decreased rapidly to less than 1 kb in alfalfa using candidate gene–related SNP (Herrmann et al. 2010). The rapid LD decrease is likely due to the outcrossing nature of alfalfa and the diverse germplasm we collected.

### The candidate genes of FD height

A total of four SNPs were determined to be associated with FD height based on the 1% Bonferroni test (Table 2 and Figure 4). Two out of four SNPs can be mapped to candidate genes based on LD information, and one SNP is connected with a DREB1C gene. DREB1C has been shown to be associated with freezing tolerance. When the gene CBF2/DREB1C was disrupted, mutant Arabidopsis had a higher capacity to tolerate freezing than WT plants and were more tolerant to dehydration and salt stress (Novillo et al. 2004). Further validation has been done in model plant Medicago truncatula. Transgenic Medicago truncatula exhibited suppressed shoot growth and enhanced freezing tolerance (Chen et al. 2010). The transcriptomic analyses of different dormancy alfalfa cultivars showed that DREB1C was one of the most significantly differentially expressed genes and was upregulated in a fall dormant alfalfa cultivar (Liu et al. 2019). Another transcriptome sequencing analysis revealed that nine DREB unigenes responded to cold and/or freezing stresses (Shu et al. 2017). FD height can be considered a signal of fall dormancy and freezing tolerance. If an alfalfa cultivar has a higher FD height, it will also have less freezing tolerance (Sheaffer et al. 1992). We further confirmed that DREB1C contributes to FD using the GWAS method. Combining several previous results from transgenic studies, transcriptomics, and GWAS showed that the DREB1C gene is very important for alfalfa freezing tolerance. Validation of the DREB1C gene effects in alfalfa using gene-modification tools such as CRISPR/CAS9 (Chen et al. 2020) or a transgenic method (Chao et al. 2014) will give us a better understanding of the function of DREB1C.

In addition to the DREB1C gene, there should be other genes responsible for FD, because FD height is a quantitative trait with continual variance (Figure 1). However, the strict standards (1% Bonferroni test and 36 kb candidate gene interval) used in this study resulted in the discovery of only two candidate genes, one of which a clear relationship with FD. There are some candidate genes responsible for FD that have been identified using omics technologies, such as ABF4 from RNA-seq results (Liu et al. 2019) and MsThi from the iTRAQ-based method (Du et al. 2018). Cross-validation from multiple omics technologies could help us develop a more accurate understanding of FD. Our results showed that GWAS results could agree with RNA-seq data. Combining GWAS methods with other omics technologies on a bigger scale could be a possible way to uncover more details regarding FD for alfalfa (Scossa et al. 2021).

In conclusion, we conducted a GWAS analysis of FD in alfalfa using whole-genome sequencing on seeding plants. Weak population structure was found among different accessions belonging to different geographical origins. Four SNPs were shown to be associated with FD. One candidate gene, DREB1C, was responsible for FD. This research will help us achieve a better understanding of the relationship between genotype and the FD phenotype. The results could serve as basic information for QTL mapping or candidate gene cloning to understand the mechanism of FD in alfalfa. Furthermore, analyzing the genetic basis of FD will help to develop cultivars with different FD.

## Materials and methods

### Plant materials and phenotyping

The plant materials used in this study consisted of 220 accessions collected from all over the world. These accessions include 55 cultivars, 26 cultivated materials, 95 landraces, 4 breeding materials, 16 wild materials, and 23 uncertain improvement status materials. Among them, 26 accessions come from the database of the Medium Term Library of National Grass Seed Resources of China. Another 194 accessions come from the database of the U.S. National Plant Germplasm System (USDA GRIN). The geographic sources of the accessions include China, the United States, Turkey, Afghanistan, Tashkent Uzbekistan, Spain, Russia, Morocco, France, Argentina, and others.

In October 2017, the seeds of 220 accessions were planted in the greenhouse of the Chinese Academy of Agricultural Sciences (CAAS) in Langfang, Hebei Province, China. Individuals were transplanted to the field of CAAS in April 2018. A randomized complete block design with three replications was used for the experimental design. Every accession included 5 individuals placed 30 cm apart in a single row for one replication. The accession spacing was 65 cm between rows and columns. To keep all individuals uniform, they were clipped to a height of 5 cm after transplant. Furthermore, all individuals were clipped again at the early flowering stage (when 10% of plants begin flowering). Plants were clipped three times in 2018 and four times in 2019. Fall dormancy height was collected using plant regrowth one month after final clipping (31 Oct 2018 and 29 Oct 2019). Yield-related traits (yield, plant height, flowering time, regrowth, spring vigor, leaf length, and leaf width) were collected in 2020. The trait of spring vigor is evaluated using a subjective score (1=small, 2=medium, 3=large) in the spring of 2020 (21 Apr 2020). The phenotypes of leaf length and leaf width were measured before flowering, after leaf size had stopped changing (25 Apr 2020). Three leaves were selected from the bottom part of each individual. Leaf length was the longest part of one leaf. Leaf width was the widest part of one leaf. The mean value of different leaves was considered as the leaf length and leaf width of one individual. The traits of the flowering time were collected from 27 Apr 2020 to 31 May 2020 when the plant generated its first flower. The traits of yield and plant height in the first cut were evaluated after flowering time traits were collected (change from 27 Apr 2020 to 31 May 2020). The yield and plant height in the second and third cut were collected when 10% of individuals began flowering (1 Jul 2020 and 4 Aug 2020). The yield was the fresh biomass yield of one individual, and plant height was the height of the highest branch of one individual. Regrowth was the plant height two weeks after the second cut (15 Jul 2020). All seven yield-related traits in 2020 were used to analyze the correlation coefficient with fall dormancy height in 2019. The distribution and correlation of phenotypes were analyzed using the R package PerformanceAnalytics (Peterson et al. 2018).

The variance of fall dormancy height was analyzed using a generalized linear model (GLM) as follows:

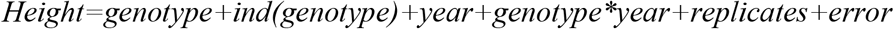

All factors were random, and the GLM was performed using PROC GLM (SAS Institute, 2010). The broad-sense heritability (H2) was computed as follows:

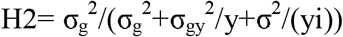

where σ_g_^2^ is the variance of genotype; σ_gy_^2^ is the interaction variance of genotype and year; σ^2^ is the residual variance. The items y and i in the equation refer to the number of years and individuals, respectively.

### Sequencing and SNP calling

Phenotyping was conducted one 15 individuals per accession and one individual with the typical phenotype was selected for sequencing. Young leaves were selected at the early regrowth stage (9 July 2019). Total DNA was extracted using the CWBIO Plant Genomic DNA Kit (CoWin Biosciences, Beijing, China), according to the manufacturer’s protocol. At least 6 ug of genomic DNA from each accession was used to construct a sequencing library following the manufacturer’s instructions (Illumina Inc.). Paired-end sequencing libraries with an insert size of approximately 300 bp were sequenced on an Illumina NovaSeq 6000 sequencer at BerryGenomics company. The data size of every accession is 10 Gb and the average Q30 is 85%. The raw data has been uploaded to the National Genomics Data Center (NGDC, https://bigd.big.ac.cn/) under BioProject PRJCA004024. Sequencing data were first quality filtered using Trimmomatic software with default parameters (Bolger et al. 2014). Paired-end sequencing reads were mapped to the alfalfa reference genome (haploid genome with 8 chromosomes (Long et al. 2021)) with BWA-MEM using default parameters (Li 2013). SAMtools were used to translate SAM file to BAM file and sort BAM files using default parameters (Li et al. 2009).

Picard Tools was used to mark duplicate reads (http://broadinstitute.github.io/picard/), and Genome Analysis ToolKit was used to correct indels which can be mistaken for SNPs (Van der Auwera et al. 2013). SAMtools mpileup and VarScan were used to detect SNPs (Koboldt et al. 2012). Furthermore, SNP data were filtered using VCFtools (Danecek et al. 2011) with a missing rate of less than 10%, a minor allele frequency of more than 0.05, and mean read depth greater than 20. The SNP distribution on the chromosome was generated using the R code from a GWAS study in beef cattle (Zhou et al. 2019).

### Population structure and linkage disequilibrium

Population structure was calculated using the TASSEL 5 software with principal component analyses (PCA) (Bradbury et al. 2007). The PCA results combined with geography information were plotted using R packages ggplot2 (Wickham 2011) and scatterplot3d (Ligges and Mächler 2002). LD information was calculated using the software PopLDdecay with default parameters (Zhang et al. 2019). The VCF file containing information on all 875,023 SNP markers was imported to PopLDdecay. The LD results among all accessions were used to estimate the LD of alfalfa. The distance versus mean R^2 within 30 kb of LD was plotted using R package ggplot2.

### Association mapping

The Bayesian-information and linkage-disequilibrium iteratively nested keyway (BLINK) method was used to carry out the GWAS (Huang et al. 2019) for mean value of fall dormancy height (FD height). GWAS was conducted using the BLINK C version software. The Bonferroni multiple test correction was used to determine the significant SNP threshold (P=0.01/875,023= 1.14×10-8). The BLINK method uses iterations to select a set of markers associated with a trait. These associated markers are fitted as a covariate for testing the remaining markers. This method better controls false positives than the kinship approach. The real data and simulated data results showed that BLINK has higher statistical power than other methods, such as MLM, FarmCPU, et al. (Huang et al. 2019). The Manhattan and Q-Q plots of GWAS results were created using the R package GAPIT3 (Wang and Zhang 2020).

## Declarations

### Ethics approval and consent to participate

Not applicable.

### Consent for publication

Not applicable.

### Availability of data and materials

All RAD raw sequence data were upload to the National Genomics Data Center (NGDC, https://bigd.big.ac.cn/) under BioProject PRJCA004024. The datasets used and analyzed during the current study are available from the corresponding author on reasonable request.

### Competing interests

The authors declare that they have no conflicts of interest.

### Funding

This work was supported by the National Natural Science Foundation of China (No. 31971758), and the breeding forage and grain legumes to increase China’s and EU’s protein self-sufficiency, collaborative research key project between China and EU (2017YFE0111000/EUCLEG 727312). The funding body played no role in the design of the study, the collection, analysis, and interpretation of data and the writing of the manuscript.

### Author contributions

QCY conceived and designed the experiments. FZ, JMK, RCL, MNL, YS, ZW, and ZWZ performed the experiments. FZ, and ZWZ analyzed the data. FZ, ZWZ, and QCY wrote the paper. All authors read and approved the final manuscript.

## Acknowledgements

We thank the database of the Medium Term Library of National Grass Seed Resources of China and the database of the U.S. National Plant Germplasm System (USDA GRIN) for providing alfalfa accessions. We thank China Scholarship Council (CSC) for their support the international study of Fan Zhang.

---

**Supplemental Figure 1.**
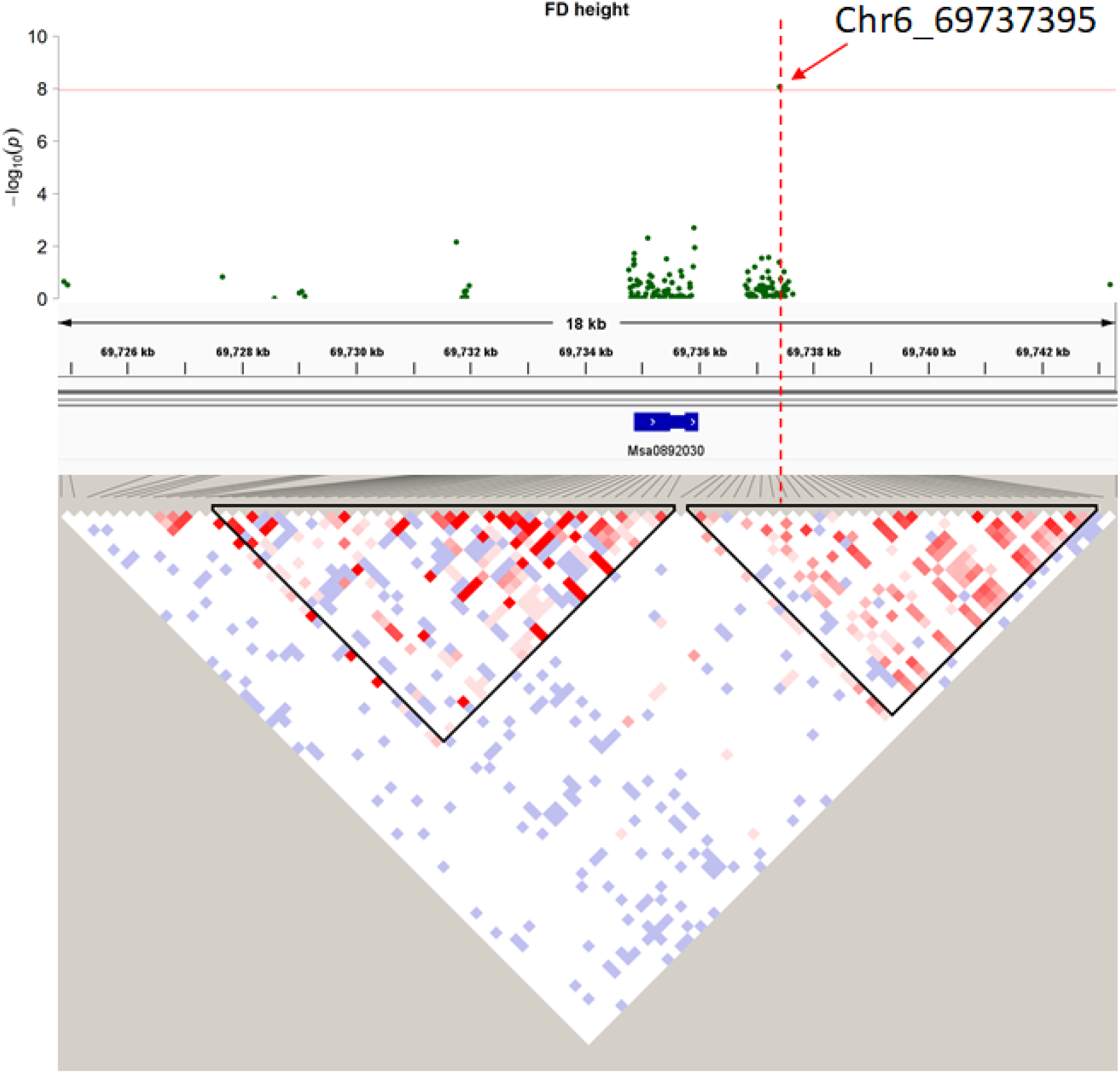
The GWAS results combined with gene information and LD information in the 36 kb interval. The Manhattan plot (top) shows the GWAS results in the 18 kb interval (chr6:69724759-69743282) belonging to the 36 kb interval SNP. Below this plot is shown the gene distribution and location in this interval. The bottom plot shows the linkage distribution results for the SNP markers in this interval. Different colors represent different LD values. The red color means a high LD value. The black triangles represent potential LD blocks.

**Supplemental Figure 2.**
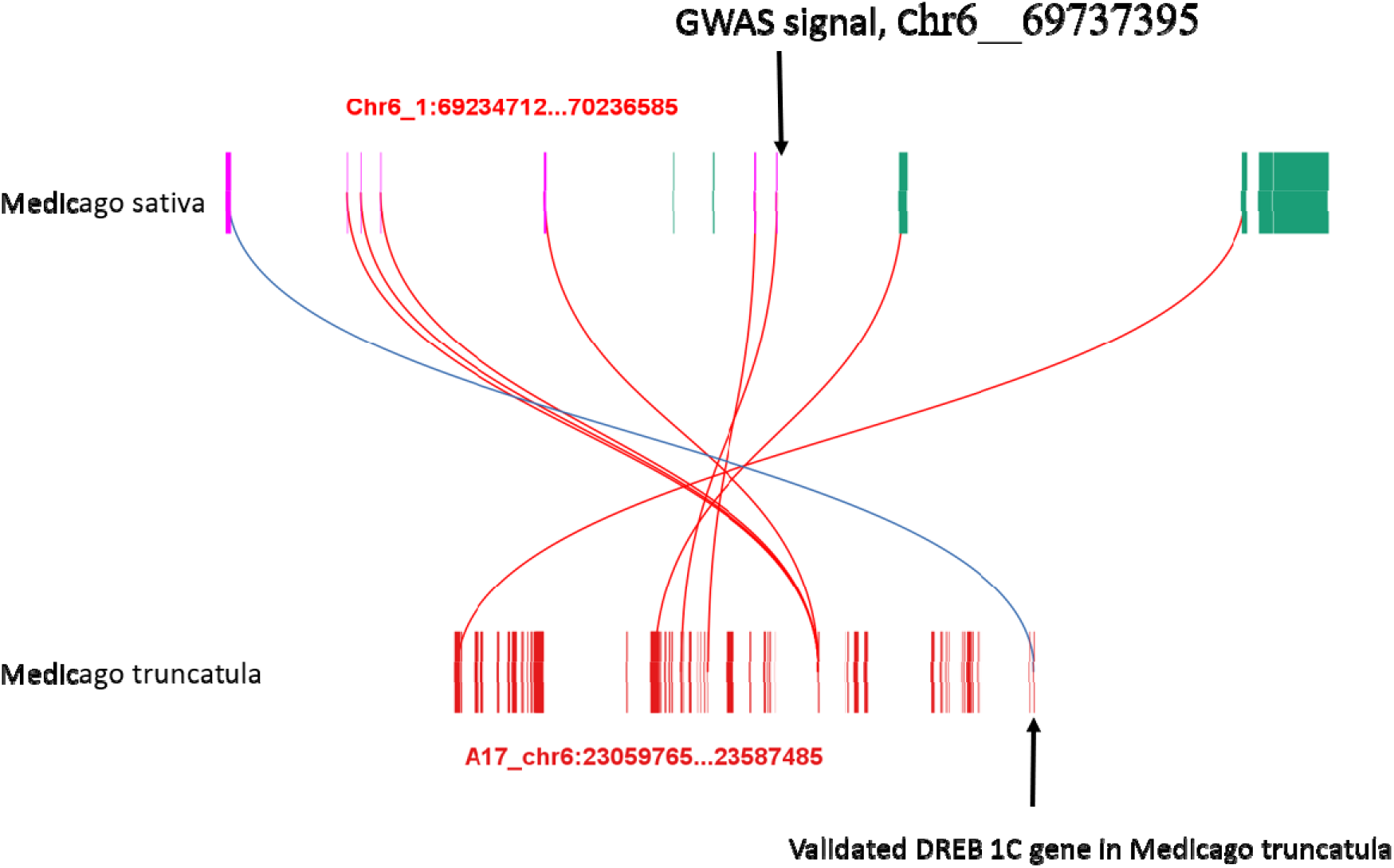
The collinearity of DREB genes in alfalfa and M. truncatula genomes. Alignment of Medicago sativa with Medicago truncatula genomes in the DREB genes region. The region 500 kb up- and downstream of the GWAS signal including 7 DREB genes is shown. The pink vertical lines represent 7 DREB genes. The green color represents other genes in this interval. The red vertical lines at the bottom of the figure are the genes of Medicago truncatula. The curved lines connecting Medicago sativa and Medicago truncatula indicate corresponding homologous genes. The alfalfa homologous DREB1C gene validated in Medicago truncatula is highlighted with the blue curve. The figure was generated using the TBtools software.

**Supplemental Table 1.** The physical position of 9 DREB genes on chromosome 6.

| Gene Name | Chr | Start_pos | Stop_pos | Gene Annotation |
| --- | --- | --- | --- | --- |
| Msa0891950 | chr6 | 69234712 | 69239739 | DREB1C |
| Msa0891960 | chr6 | 69344692 | 69345384 | DREB1C |
| Msa0891970 | chr6 | 69357058 | 69357738 | DREB1C |
| Msa0891980 | chr6 | 69374816 | 69375502 | DREB1C |
| Msa0891990 | chr6 | 69523664 | 69525980 | DREB1C |
| Msa0892020 | chr6 | 69715010 | 69716598 | DREB1C |
| Msa0892030 | chr6 | 69734863 | 69735980 | DREB1C |
| Msa0893350 | chr6 | 76949157 | 76950761 | DREB1C |
| Msa0893360 | chr6 | 76953910 | 76957350 | DREB1C |

## Notes

https://bigd.big.ac.cn/

http://broadinstitute.github.io/picard/

### Competing Interest Statement

The authors have declared no competing interest.

